# Neurocomputational Mechanisms Linking Future Interaction Prospects to Reactive and Proactive Costly Punishment

**DOI:** 10.1101/2025.05.15.654171

**Authors:** Chuangbing Huang, Xingmei Zhou, Feilong Liu, Binjie Yang, Zixin Zheng, Xiaoqing Li, Yijie Yin, Yue-Jia Luo, Chunliang Feng

## Abstract

Costly punishment within social bonds manifests as both reactive penalties for transgressions and proactive sanctions against fair behaviors, yet a unified neurocomputational framework remains elusive. Combining neuroimaging and computational modeling, this study examined how future interaction prospects—a key element of social bonds—modulate reactive and proactive punishment. Participants acted as second- or third-party punishers, responding to allocation offers from individuals with or without future interactions. Results showed that participants imposed harsher punishment on both fair and unfair offers from individuals they did not expect to interact with, aligning with lower self-reported social closeness. Heightened provocation sensitivity, reflected in dorsal anterior cingulate cortex activity, drove the increased reactive punishment toward wrongdoers without future interactions, whereas punishment of future-interacting transgressors was associated with enhanced dorsolateral and dorsomedial prefrontal cortex (dlPFC, dmPFC) activity. A stronger preference for relative advantage and reduced harm sensitivity, linked to dlPFC and temporoparietal junction activity, mediated greater proactive punishment towards non-future-interacting individuals. Moreover, proactive punishment and underlying neurocognitive processes followed a collective retaliation pattern, increasing after unfair (vs. fair) behavior from another non-future-interacting individual. These findings establish a unified neurocomputational framework for understanding how future interaction prospects regulate costly punishment, integrating reactive and proactive motives within social bonds.

## Introduction

Costly punishment, which entails incurring a personal cost to sanction others, plays a critical role in norm enforcement and cooperation among genetically unrelated individuals. As a hallmark of human societies, it has attracted substantial attention across psychology, economics, and evolutionary biology [1–4]. However, beyond norm enforcement, costly punishment may engage a multitude of additional motives within social bonds where most social interactions are embedded. For instance, the same norm violations (e.g., unfair allocations) elicit different punitive responses depending on the punisher’s social ties with the transgressor—such as shared nationality, preferences, friendship, or romantic ties—highlighting the role of affective preferences [5–8]. Furthermore, costly punishment is not limited to sanctioning transgressions; it can also target norm-abiding behaviors, manifesting as proactive (or ‘antisocial’) punishment [9]. This form of punishment is more frequently directed at distant others [8, 10, 11] and may be driven by motives such as collective retaliation and status-seeking. Despite their significance, how these motives jointly shape costly punishment within social bonds remains largely unexplored. This study addressed this gap by examining neurocomputational mechanisms through which the prospect of future interactions—an essential feature of social bonds—modulates both reactive and proactive costly punishment.

Existing literature presents mixed findings on how social bonds influence reactive punishment [12]. While some studies suggest that individuals are more lenient toward norm violations by close affiliates than by distant others [7, 13–15], others report the opposite [16–20]. The disparate findings likely reflect the multidimensional nature of social bonds, which involve factors such as similarity, familiarity, and functional interdependence, with different dimensions being more relevant in different contexts [12, 21]. Thus, simply determining whether reactive punishment is influenced by social bonds provides limited insight, as social bonds vary across multiple dimensions. The current study tackled this issue by isolating and assessing the impact of future interaction prospects. Future interaction expectations represent a defining feature of virtually all social bonds, including those formed arbitrarily within experimental settings. For instance, future interaction expectations mediate the effect of arbitrary group assignments on preferences toward in-group members [22, 23]. Conversely, shifting these expectations from in-group to out-group members reduces in-group favoritism [24].

Moreover, the prospect of future interaction is also well-suited to disentangle two opposing perspectives on the relationship between social bonds and reactive punishment [12]. According to the norm enforcement hypothesis, punishment fosters cooperation within one’s social network by regulating the behavior of those with whom one expects to interact [17, 22]. By this account, reactive punishment will be disproportionately directed toward potential future partners. In contrast, the mere preference hypothesis suggests that social closeness mitigates moral outrage over norm violations [12]. Therefore, anticipating future interactions will reduce the likelihood of reactive punishment, as this prospect promotes interdependence and affective preferences [24]. While these accounts make opposite predictions, both posit that social bonds influence punishment by altering sensitivity to provocation or economic self-interest.

Supporting this view, social bonds modulate activity in brain regions associated with encoding aversive feelings elicited by norm violations, including the anterior insula (AI) and dorsal anterior cingulate cortex (dACC) [8, 25, 26], as well as regions involved in inferring violators’ intentions, such as the temporoparietal junction (TPJ) and medial prefrontal cortex (mPFC) [25, 27]. Additionally, social bonds influence activity in the dorsolateral prefrontal cortex (dlPFC), a key node of the deliberate system implicated in integrating provocation signals and self-interest computations to facilitate flexible and strategic punishment [8, 28]. While prior studies consistently implicate these regions, the specific computations they implement remain largely unknown. Furthermore, whether expectations of future interactions engage similar neural mechanisms requires further investigation.

Despite extensive research on reactive punishment, proactive punishment within social bonds remains understudied. Emerging evidence indicates that individuals punish fair behaviors more harshly for distant others than affiliated others [8, 10]. Such behaviors cannot be reconciled with norm enforcement motives, implying alternative mechanisms driven by affective preferences. One possibility, informed by intergroup categorization research, is that people are more inclined to classify distant others into broad social categories, perceiving them as interchangeable [29]. Thus, a recent norm violation by one distant individual may elicit proactive punishment to uninvolved others within the same category, reflecting collective retaliation [30]. Collective retaliation seeks to impose harm and often involves attributing responsibility to innocent individuals based on their social ties to actual wrongdoers [31], a process closely linked to the mentalizing network (e.g., TPJ). Alternatively, self-enhancement research suggests that individuals view themselves and close associates as superior to distant others and engage in behaviors reinforcing this belief [32]. Proactive punishment may thus serve status-seeking motives aimed at securing a relative advantage [33, 34]. As such, proactive punishment primarily reflects strategic considerations mediated by the deliberative system (e.g., dlPFC). However, empirical support for these mechanisms remains scarce. This study is the first to examine whether future interaction expectations shape proactive punishment by modulating sensitivity to harming others and/or concerns for relative advantage, as well as their underlying neurobiological basis.

By isolating the role of future interaction prospects and integrating computational modeling with neuroimaging, this study aimed to advance a unified framework for understanding how social bonds shape costly punishment. Specifically, we dissected the distinct neurocomputational processes through which future interaction expectations influence reactive and proactive punishment, including sensitivity to provocation, economic self-interest, harm, and relative advantage.

## Methods

### Participants

Thirty Chinese healthy college students (17 males, mean age ± S.D. = 20.20 ± 1.64 years) participated in the study for monetary compensation. The sample size was based on previous fMRI studies on the similar topics [3, 25, 35, 36] and resource constraints. All participants were right-handed, had normal or corrected-to-normal vision, and had no history of neurological or psychiatric disorders. Written informed consent was obtained from all participants. The study was conducted according to the ethical guidelines and principles of the Declaration of Helsinki and was approved by the local ethical committee.

### Experimental procedure

Participants played second- and third-party punishment games during fMRI scanning. In the second-party punishment (SPP) game (**Fig. 1A**), there were two players: a dictator and a recipient, with the participants taking on the recipient role. The dictator distributed a sum of money (12 monetary units, MUs) between two players, and the recipient could only accept the allocation [37]. For each trial, both the participant (as recipient) and the dictator were given an additional 6 MUs. In response to each dictator’s division of money, the recipient decided how many MUs to spend to punish the dictator by decreasing dictator’s payoffs: each MU spent by the recipient reduced the payoff of the dictator by 3 MUs [1, 38, 39]. The third-party punishment (TPP) game (**Fig. 1B**) was similar to SPP game, except that the participants acted as an agent on behalf of another recipient, thereby making punishment decisions that affected the outcomes for the recipient and the dictator [40]. The current study employed both SPP and TPP games to assess the robustness of future interaction effects on costly punishment, as the extent to which these paradigms engage similar cognitive subcomponents remains debated [41, 42]. Importantly, words such as “fairness,” “punishment,” “sanction,” or “dictator” were not used in the instructions, and participants were told that they could conduct “deduction money” to dictators [1]. To encourage real decisions from participants, it was emphasized that MUs were convertible to monetary payoff, and that all players would be paid according to their choices in the game, in addition to a fixed show-up compensation.

**Fig. 1.**
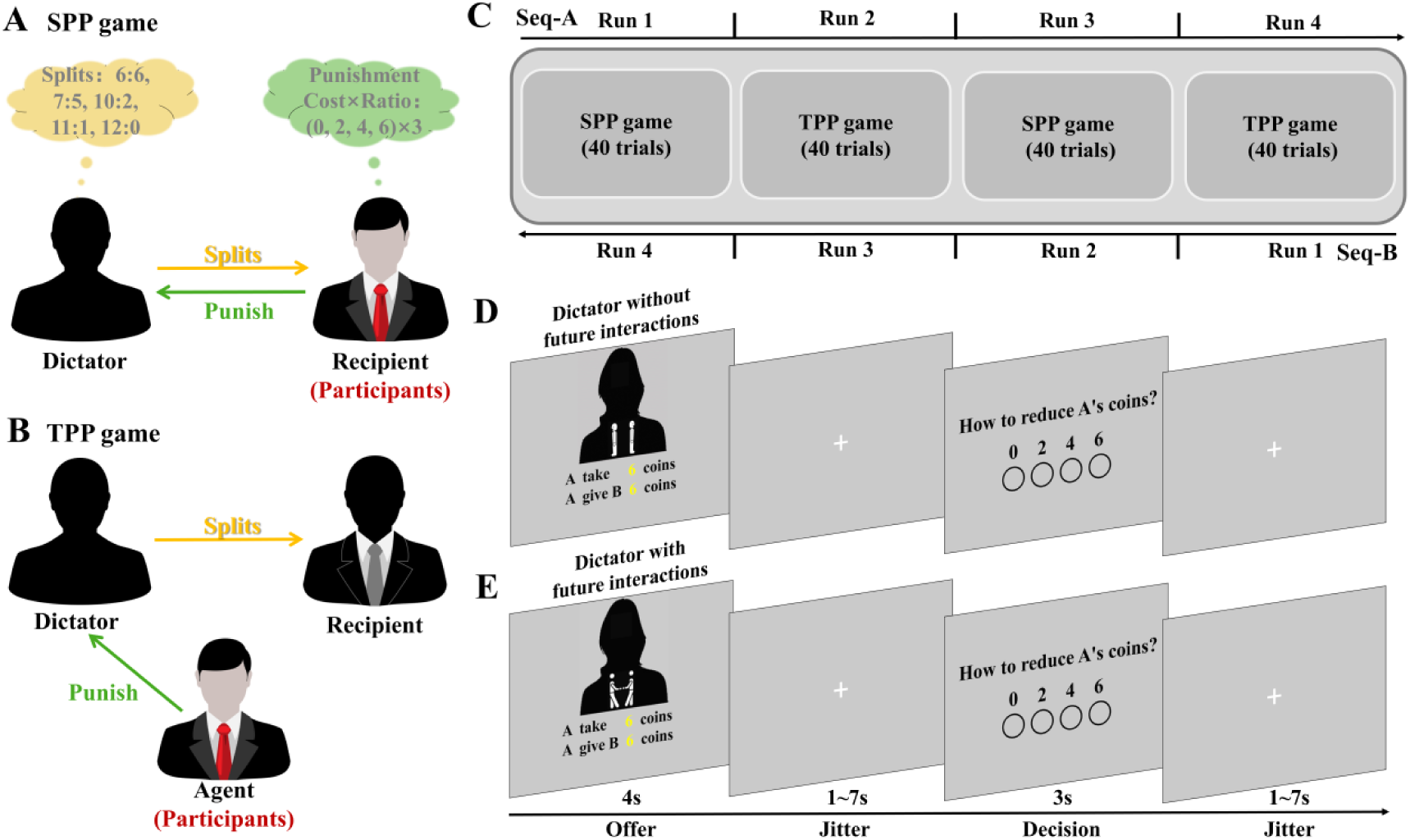
Task design and experimental procedure. **(A)** Participants as the second-party received splits from dictators and decided how to punish the dictators. **(B)** Participants as the third-party decided how to punish the dictators on behalf of another recipient. **(C)** The assignment of SPP and TPP sequences was counterbalanced across participants. **(D)** The timeline of trial for dictators without future interactions. **(E)** The timeline of trial for dictators with future interactions. SPP, second-party punishment; TPP, third-party punishment; Seq, sequence.

Participants encountered dictators either with or without expectations of future interactions [24, 43]. In the condition involving future interactions, participants were informed that both they and the represented recipient would interact with these dictators in a subsequent session. In the condition without future interaction, participants were told that neither they nor the represented recipient would interact with these dictators again. Across both conditions and experimental sessions, all decisions were made anonymously. However, participants were informed that dictators would receive feedback on the punitive responses made by anonymous recipients. Unbeknownst to them, all participants were assigned the roles of SPP or TPP punishers, with no actual dictators or subsequent sessions. Before leaving the laboratory, participants were debriefed, and none expressed doubts regarding the experimental procedures.

The fMRI scanning protocol consisted of four runs, with SPP and TPP presented in an alternating sequence (i.e., SPP-TPP-SPP-TPP or TPP-SPP-TPP-SPP, **Fig. 1C**). The assignment of SPP and TPP sequences was counterbalanced across participants. Before each run, participants were informed of their current role (SPP or TPP). Each run consisted of 40 trials, with 20 trials of fair splits (10 trials of 6:6 splits, 10 trials of 7:5 splits) and 20 trials of unfair splits (8 trials of 10:2 splits, 6 trials of 11:1splits, and 6 trials of 12:0 splits) for both types of interaction expectations (with versus without). Participants were informed that each allocation on every trial was proposed by a different dictator.

On each trial (**Fig. 1D and 1E**), information on future interactions and allocations were displayed for 4 s and followed by a jitter (1 ~ 7 s). Next, the decision phase was presented for 3 s, during which participants decided how many MUs to spend to reduce the dictator’s payoff by selecting one of four possible choices (a hollow circle was below each choice): 0, 2, 4 or 6 MUs. Choices were made through a response box, with associations between buttons and decisions being counterbalanced across participants. Once participants’ response was collected, the corresponding choice was highlighted with a filled circle and was displayed on screen until the end of the decision phase. Each trial ended with a second jitter (1 ~ 7 s). Stimulus presentation and behavioral data collection were conducted by using Psychtoolbox-3.

After fMRI scanning, participants completed a survey assessing social closeness with dictators across interaction expectations (with/without), splits (6:6/7:5/10:2/11:1/12:0), and roles (SPP/TPP). Closeness was rated on a 9-point Likert scale (1 = very distant, 9 = very close) in response to the question: “How close did you feel to the dictator?”

### fMRI data acquisition

Imaging was performed on a 3T SIEMENS MAGNETOM Prisma scanner equipped with a 64-channel transmit/receive gradient head coil at Shenzhen University’s Imaging Center for Brain Research. A T2-weighted gradient-echo-planar imaging (EPI) sequence was used to acquire functional images: TR = 1500 ms, TE = 30 ms, flip angle = 75°, number of axial slices = 72, slices thickness = 2.0 mm, matrix size = 96 × 96 mm^2^, and FOV = 192 × 192 mm^2^. High-resolution anatomical images covering the entire brain were obtained by applying magnetization-prepared rapid acquisition with a gradient-echo (MPRAGE) sequence: TR = 1900 ms, TE = 2.23 ms, flip angle = 7°, number of slices = 224, slices thickness = 1.10 mm, matrix size = 192 × 192 mm^2^, FOV = 220 × 220 mm^2^.

### Statistical analysis Behavioral analysis Amounts of punishment

Analyses of behavioral data were performed using SPSS 21.0 (IBM, Somers, USA) with a threshold of *p* < 0.05 (two-tailed). The amounts of punishment were analyzed with a 2 (Interaction expectation: with/without) × 2 (Split: fair/unfair) × 2 (Role: SPP/TPP) repeated-measures analysis of variance (ANOVA). **Distributions of punishment choices.** The average punishment amounts were calculated by aggregating the frequency of each punishment choice (0, 2, 4, and 6). A follow-up analysis examined how the distribution of punishment choices varied as a function of interaction expectations. For each split (i.e., 6:6, 7:5, 10:2, 11:1, 12:0), a separate Friedman rank-sum test was conducted to assess the effect of interaction expectations on the distribution of punishment choices.

### Effects of trial sequence on punishment decisions

The collective retaliation account predicts that a recent norm violation by one individual triggers proactive punishment toward others in the same category [30]. To test this hypothesis, we examined sequential trial effects on punishment, assessing whether participants’ punishment decisions in each condition were influenced by the contexts of the immediately preceding trial. Taking trials of fair splits from dictators without interaction expectations as an example, we computed average punishment amounts based on four preceding contexts: fair splits from dictators without future interactions, unfair splits from dictators without future interactions, fair splits from dictators with future interactions, and unfair splits from dictators with future interactions. Punishment amounts for each condition were then analyzed using a 2 (Preceding Interaction expectation: with/without) × 2 (Preceding Split: fair/unfair) repeated-measures ANOVA.

### Self-reported closeness

The self-reported social closeness was analyzed with a 2 (Interaction expectation: with/without) × 2 (Split: unfair/fair) × 2 (Role: SPP/TPP) repeated-measures ANOVA.

Moreover, the relationship between self-reported closeness and punishment amounts was examined using representational similarity analysis (RSA) [44]. Specifically, a 20 × 20 representational dissimilarity matrix (RDM) was separately constructed for self-reported closeness and punishment amounts, computed as Euclidean distances based on the average closeness ratings and punishment amounts across participants for each pair of conditions. These conditions were derived from a 2 (Interaction expectation: with/without) × 5 (Split: 6:6/7:5/10:2/11:1/12:0) × 2 (Role: SPP/TPP) design. The correlation between the resulting RDMs was then computed to assess the association between self-reported closeness and punishment amounts. The statistical significance of this correlation was evaluated using a permutation test with 10,000 iterations. In each iteration, the condition labels of the two RDMs were randomly shuffled, and the correlation was recalculated to generate a null distribution of correlation coefficients. The *p*-value was determined as the proportion of permuted correlation coefficients exceeding the observed correlation coefficient, divided by the total number of permutations (i.e., 10,000).

### Computational modeling

To dissociate the psychological processes underlying the effects of interaction expectations on punishment decision-making, we developed computational models based on hypotheses derived from norm enforcement and mere preference accounts [12, 25, 30]. Specifically, we modeled participants’ punishment behavior within the inequity aversion framework [45]. This widely used model assumes that costly punishment decisions are driven by concerns about both inequity and monetary payoffs. Building on this model, we incorporated additional psychological components to formally test how interaction expectations shape punishment decisions, including provocation sensitivity, relative advantage concerns, economic self-interest, harm sensitivity, and retaliation preference.

#### Inequity aversion model (M1)

The standard inequity aversion model assumes that participants, as punishers, consider both disadvantageous and advantageous inequity. The utility of choice takes the following forms (Fehr & Schmidt, 1999; Zhong et al., 2016):

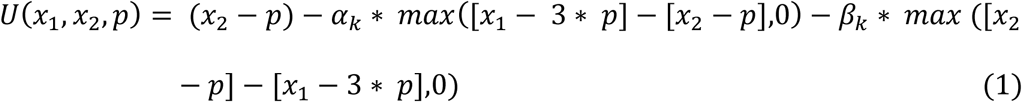

In this model, *p* represents participants’ punishment choices, *x_1_* and *x_2_* denote the payoffs of the dictator and recipient, respectively. The parameters *α_k_* (range, 0 ~ +∞) quantify sensitivity to disadvantageous inequity arising from dictators’ unfair allocations (i.e., provocations), with larger values indicating greater provocation sensitivity. The parameters *α_k_* vary across participants and conditions, where *k* = [1, 2, 3, 4] corresponds to the following conditions: dictators with interaction expectations in the SPP game, dictators with interaction expectations in the TPP game, dictators without interaction expectations in the SPP game, and dictators without interaction expectations in the TPP game. The parameters *β_k_*(range, -∞ ~ +∞) capture sensitivity to advantageous inequity, with larger values indicating stronger aversion and smaller values reflecting a preference for relative advantage [45, 46]. Advantageous inequity in this study primarily resulted from proactive punishment of fair allocations (i.e., 6:6 and 7:5), where any nonzero punishment led to relative advantage.

#### Inequity aversion plus cost aversion model (M2)

This model extends the standard inequity aversion framework by incorporating sensitivity to personal economic costs. Given that punishment decisions entail financial losses, participants may exhibit varying degrees of cost sensitivity across different contexts [25, 47], in addition to their concerns about inequity. This model thus assumes that interaction expectations impact costly punishment by modulating sensitivity to economic costs, norm violation and relative advantage. The utility function is defined as follows:

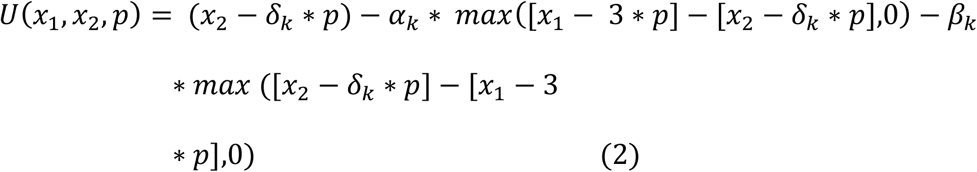

In this model, the parameters *δ_k_* (range, 0 ~ +∞) quantify sensitivity to personal costs associated with punishment across conditions, with larger values indicating stronger aversion to economic losses.

#### Inequity aversion plus harm sensitivity model (M3)

This model incorporates sensitivity to inflicted harm toward dictators. Unlike the inequity aversion plus cost aversion model, which accounts for economic self-interest, this model assumes that participants vary in their sensitivity to harming dictators across contexts [47, 48]. The utility function is defined as follows:

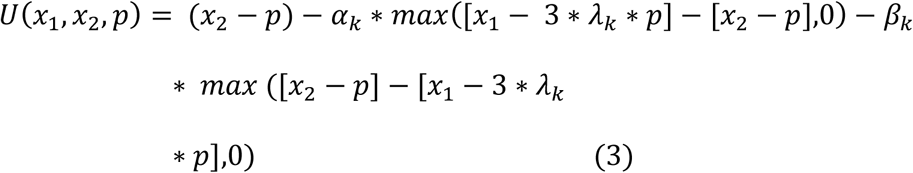

In this model, the parameters *λ_k_* (range, 0 ~ +∞) quantify sensitivity to harming dictators through punishment across conditions, with larger values indicating stronger harm aversion.

#### Inequity aversion plus retributive preference model (M4)

This model expands the standard inequity aversion framework by incorporating retributive preference, specifically, the desire to impose ‘just deserts’ on perpetrators [33]. That is, participants respond to provocation not only due to inequity concerns but also proportional retaliation. The utility function is defined as follows:

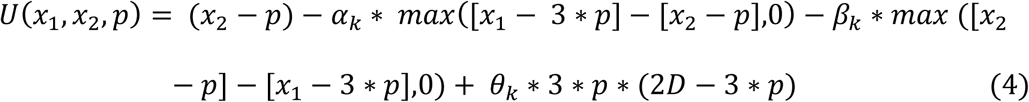

In this model, *D* represents the degree of provocation, calculated as the difference in payoffs between the dictator and recipient due to the dictator’s initial allocation. Retributive motivations are maximized when the participant’s punishment (3\**p*) closely matches the degree of provocation. The parameters *θ_k_* (range, 0 ~ +∞) quantify preferences for retaliation across conditions, with larger values indicating a stronger inclination to punish in proportion to perceived injustice.

Based on the utility of each choice, we used the softmax function to estimate the probability of each punishment choice:

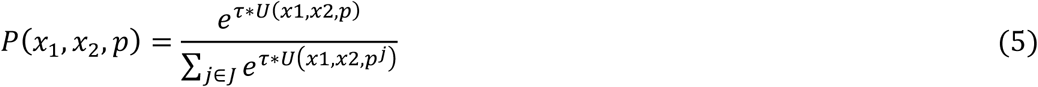

Where the parameter *τ* (range, 0 ~ +∞) is the softmax inverse temperature. The smaller *τ* is, the more randomly participants behaved, and *p^j^* denotes the possible punishment decisions that could be implemented in each trial.

### Model comparison

The leave-one-out information criterion (LOOIC) and widely applicable information criterion (WAIC) were employed as the metrics for model comparison [49]. Both LOOIC and WAIC estimate out-of-sample pointwise predictive accuracy using the posterior simulations. LOOIC approximates the accuracy through leave-one-out cross-validation, while WAIC is based on the series expansion of leave-one-out-cross-validation [49]. By convention, lower LOOIC or WAIC values indicate better prediction accuracy. The model with the lowest WAIC and LOOIC is considered as the winning model. As a general guideline, a difference of 10 points in LOOIC or WAIC between models can demonstrate the superiority [50]. LOOIC and WAIC values for all candidate models were computed using the “loo” package in R. Additionally, we employed Bayesian model averaging to calculate model likelihoods and derive LOOIC and WAIC weights. These weights represent the probability that each model is the best-fitting model given the data and the set of candidate models [51].

### Parameter estimation

Parameter estimation was conducted using hierarchical Bayesian analysis, which offers more stable and precise estimation compared to maximum likelihood estimation [52]. The hierarchical Bayesian analysis was implemented using the “RStan” package in the R programming language. The “RStan” package relies on the Stan interface [53], which employs Markov Chain Monte Carlo sampling methods to perform fully Bayesian inference and obtain posterior distributions. Following the approach in “hBayesDM” package [52], it was assumed that individual level parameters were drawn from group level normal distribution Normal (*μ_0_*, *σ_0_*) where *μ_0_* and *σ_0_* refer to group-level mean and standard deviation, in other words, the hyper parameters of individual-level parameters. Weakly informative priors were adopted for the priors of the group-level normal means and standard deviations following a recommanded appraoch [52]: *μ_0_* ~ Normal (0, 10) for parameters with unconstrainted ranges, *μ_0_* ~ Normal (0, 1) for parameter with constrainted ranges, and *σ_0_* ~ half-Cauchy (0, 2) for alll parameters. This was to minimize the impact of priors on the posterior distributions. The model and priors were defined in a Stan file, which was then compiled in the R environment. Each model was fitted with 3 chains, each consisting of 4000 iterations after a warmup period of 2000 iterations. Based on the estimated parameters of the winning model, the posterior highest density interval (HDI) represented the uncertainty in the estimated parameters. Meaningful differences were determined if 95% HDI of the posterior was above or below 0 [52].

### Model validation

Model validation was conducted to further assess the performance of the winning model. First, we examined the correlation between the average punishment amounts in actual behavioral data and model predictions across 160 trials. Second, we correlated the average punishment amounts of actual behavioral data and model predictions across 30 participants. Third, the simulated data underwent the same statistical analyses used on the real data to determine whether the simulated data successfully recovered the behavioral effects observed in the original dataset.

### Parameter recovery

We ran parameter recovery analyses to assess how accurate a model estimates true parameter values from the simulated choice data generated from the true parameter values [54, 55]. For this purpose, the estimated posterior means from actual data were used as the true parameter values, which provided plausible combinations of parameter values for parameter recovery analysis [56]. Afterward, simulated choice data were generated by using the true parameter values (for 30 participants and 160 trials per participant). Finally, we used the simulated choice data to estimate the parameters, and then evaluated the model performance by calculating correlations between the true and predicted parameter values.

### fMRI data analysis

Functional neuroimaging data were analyzed using SPM12. Preprocessing included realignment (rigid-body registration for head motion correction), slice timing correction, normalization to Montreal Neurological Institute (MNI) space, and spatial smoothing (FWHM = 6 mm). Subsequently, several general linear models (GLMs) were constructed to identify neural signatures associated with the effects of interaction expectations on punishment behavior and related cognitive subcomponents.

The GLM1 was designed to identify brain regions involved in the impact of interaction expectations on reactive and proactive punishment, specifically examining the Interaction expectation (with/without) × Split (fair/unfair) interaction at the split phase. This interaction was of interest based on behavioral findings demonstrating its influence on punishment behavior across roles (see Results). For each participant, boxcar regressors were defined for each epoch of the time course. The split phase (4 s) and decision phase (3 s) were modeled separately, with four regressors per phase, corresponding to the Interaction expectation × Split conditions for each run of second-/third-party punishment. These regressors were convolved with the canonical hemodynamic response function and incorporated into the design matrix along with six head movement parameters. Temporal autocorrelations were corrected using a first-order autoregressive model.

The GLM2 was constructed to examine neural signatures underlying the effects of trial sequence on proactive punishment, specifically focusing on trials of fair splits from dictators without future interactions, where behavioral effects of trial sequence were identified (see Results). Four regressors were defined for both the split and decision phases, based on the preceding trial condition: fair splits from dictators without future interactions, unfair splits from dictators without future interactions, fair splits from dictators with future interactions, and unfair splits from dictators with future interactions. An additional regressor accounted for the other three conditions of non-interest, including unfair splits from dictators without future interactions, fair splits from dictators with future interactions, and unfair splits from dictators with future interactions. At the second level, we investigated the effects of trial sequence, specifically examining the interaction between Preceding Interaction expectation (with/without) and Preceding Split (fair/unfair) during the split phase.

The GLM3 was utilized to identify brain regions encoding the subjective utility of the chosen option, as determined by the winning model. The GLM3 included two regressors: one corresponding to the split phase and another to the decision phase with subjective utility as a parametric modulator. At the second level, we estimated the parametric modulations of subjective utility using a one-sample t-test.

For false positive control, we used whole-brain cluster correction with a cluster-defining threshold of *p* < 0.001 and a Family Wise Error (FWE) corrected threshold of *p* < 0.05 unless otherwise stated [57, 58].

### Whole-brain Regression analysis

Based on the winning model (the inequity aversion plus harm sensitivity model, M3, see Results), we examined the neural mechanisms underlying sensitivity to provocation (*α*), relative advantage (*β*), and inflicted harm (*λ*) during punishment decisions. Participant-specific parameter estimates from the winning model were included as covariates at the second level [3, 59]. The differences in changes of provocation sensitivity (*α*) between interaction expectations were included in the analysis of the contrast between interaction expectations in unfair splits (i.e., unfair splits from dictators with future interactions vs. unfair splits from dictators without future interactions). Moreover, the difference values of relative advantage sensitivity (*β*) were included in the analysis of the contrast between interaction expectations in fair splits (i.e., fair splits from dictators with future interactions vs. fair splits from dictators without future interactions). Lastly, the difference values of harm sensitivity (*λ*) were included in the analysis of the contrast between interaction expectations in both fair and unfair splits. All regression analyses focused on the decision phase, aiming to disentangle neural signatures associated with different component processes of subjective utility.

## Results

### Participants imposed harsher punishment on dictators without future interactions for both fair and unfair splits

The ANOVA on punishment amounts revealed significant main effects of Split (*F_1,29_* = 244.017, *p* = 1.174 × 10^-15^, *η^2^* = 0.894) and Interaction expectation (*F* = 26.437, *p* = 1.700 × 10^−5^, *η^2^* = 0.477), indicating that participants punished unfair splits more than fair splits and dictators without future interactions more than those with future interactions. In addition, the interaction of Split × Interaction expectation was significant (*F_1,29_*= 8.033, *p* = 8.277 × 10^−3^, *η^2^* = 0.217, **Fig. 2A**). Post-hoc comparisons indicated that participants punished dictators without future interactions more harshly in both fair splits (*p* = 1.620 × 10^−4^) and unfair splits (*p* = 7.300 × 10^−5^).

**Fig. 2.**
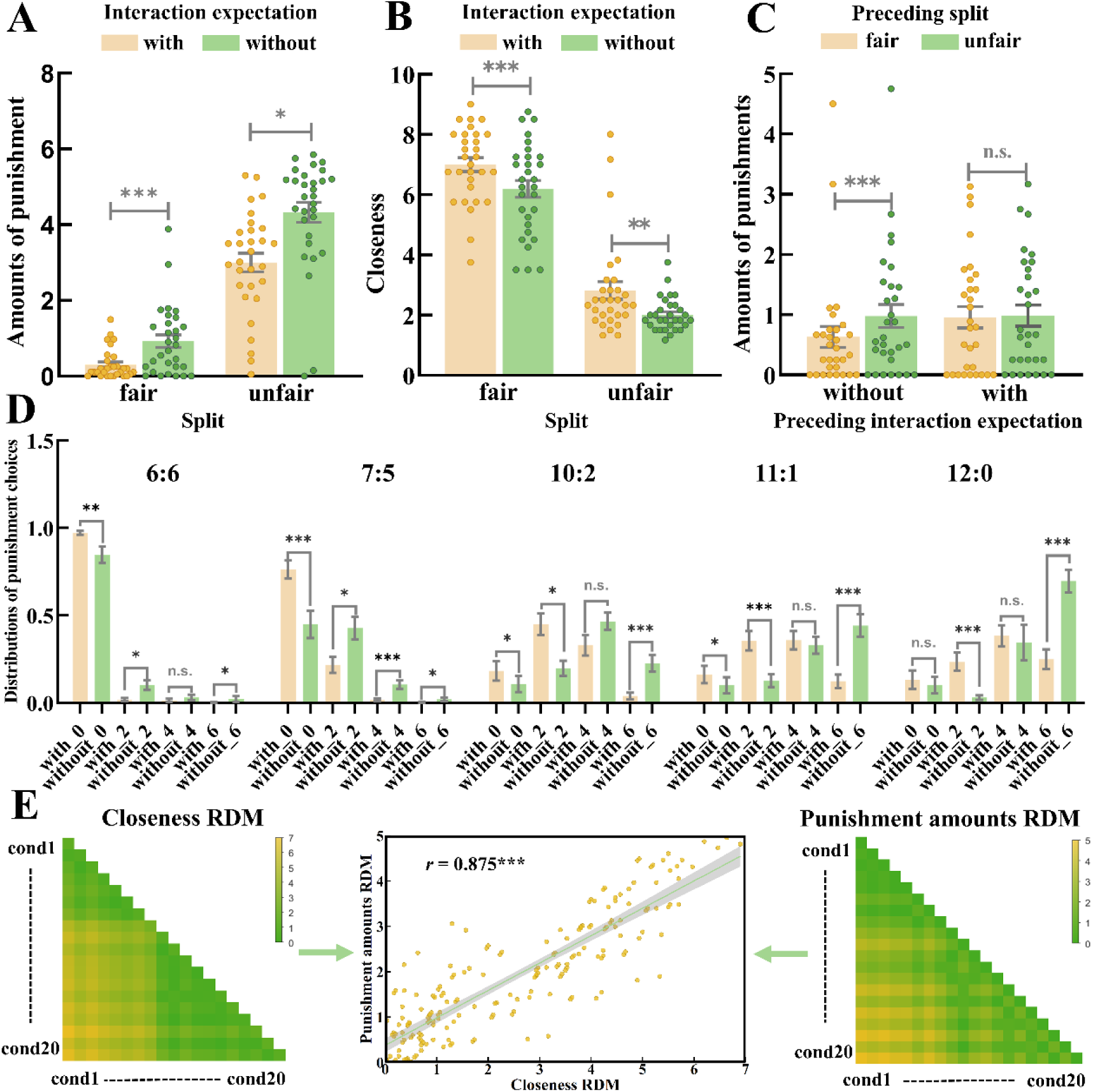
Behavioral results. **(A)** Amounts of punishment. Participants imposed harsher punishment to dictators without future interactions than those with future interactions for both fair and unfair splits. **(B)** Self-reported closeness. Participants reported higher social closeness to fair dictators and dictators with future interactions. **(C)** Trial sequence effects on punishment to fair splits of dictators without future interactions. Participants increased punishment to fair splits of dictators without future interactions when preceded by unfair than fair splits of dictators without future interactions. **(D)** Distributions of punishment choices as a function of social context and splits. **(E)** RSA analysis on the association between self-reported closeness and punishment amounts. Error bars indicate standard error. With, with future interactions; Without, without future interactions; RDM, representational dissimilarity matrix; \**p* < 0.05; \*\**p* < 0.01; \*\*\**p* < 0.001; n.s., not significant.

To further examine the modulations of Interaction expectation on the distribution of punishment choices, we conducted Friedman rank-sum tests for each split (6:6, 7:5, 10:2, 11:1, and 12:0). For fair splits (**Fig. 2D**), participants were more likely to impose 2 MUs, 4 MUs or 6 MUs on dictators without future interactions (split of 6:6, *p_2 MUs_* = 1.241 × 10^−2^; *p_6 MUs_* = 4.550 × 10^−2^; split of 7:5, *p_2 MUs_* = 1.233 × 10^−2^; *p_4 MUs_* = 2.750 × 10^−4^; *p_6 MUs_* = 2.534 × 10^−2^), but more likely to choose 0 MU for dictators with future interactions (split of 6:6, *p_0 MUs_*= 2.700 × 10^−3^; split of 7:5, *p_0 MUs_* = 1.069 × 10^−3^). For unfair splits (**Fig. 2D**), participants were more likely to choose 6 MUs to punish dictators without future interactions (split of 10:2, *p_6 MUs_* = 5.790 × 10^−4^; split of 11:1, *p_6 MUs_* = 1.200 × 10^−5^; split of 12:0, *p_6 MUs_*= 2.700 × 10^−5^), while opting for 0 or 2 MUs against dictators with future interactions (split of 10:2, *p_0 MUs_* = 2.013 × 10^−2^; *p_2 MUs_* = 2.334 × 10^−2^; split of 11:1, *p_0 MUs_* = 4.550 × 10^−2^; *p_2 MUs_* = 1.091 × 10^−3^; split of 12:0, *p_2 MUs_* = 1.300 × 10^−5^). These findings confirmed that participants punished fair and unfair splits from dictators without future interactions more severely.

In line with punishment decisions, our results also demonstrated the modulations of Interaction expectation on self-reported closeness (**Fig. 2B)**, revealing significant main effects of Interaction expectation (*F_1,29_* = 20.484, *p* = 9.500 × 10^−5^, *η^2^* = 0.414, **Fig. 3A**) and Split (*F_1,29_* = 189.687, *p* = 2.972 × 10^−14^, *η^2^* = 0.867). Participants felt closer to dictators with future interactions than without, and to dictators with fair allocations than unfair ones.

**Fig. 3.**
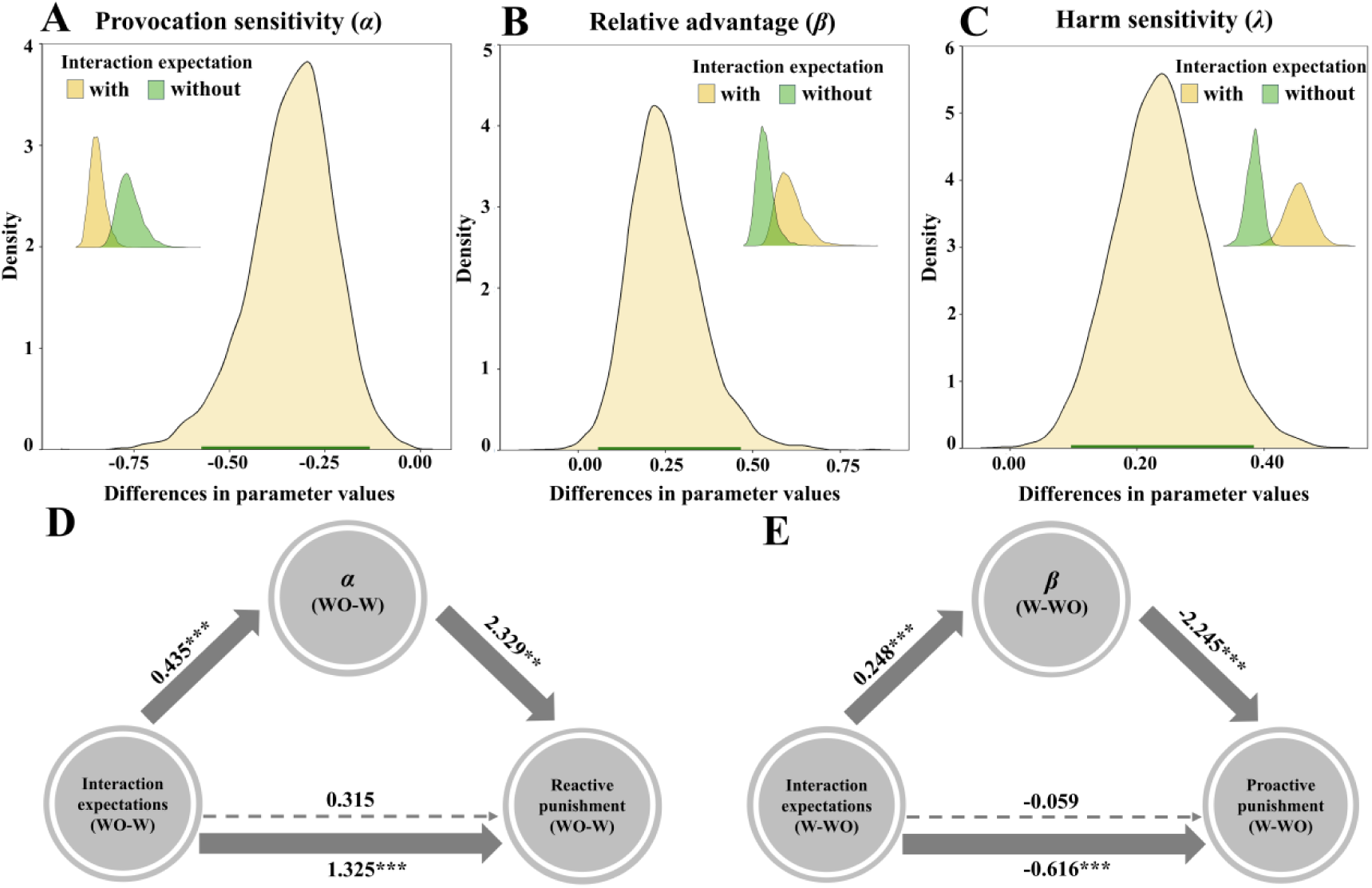
Parameters estimation and mediation results. **(A)** Sensitivity to provocation (*α*) was higher for dictators without future interactions than those with future interactions. **(B)** Relative advantage (*β*) preference was higher for dictators without future interactions than those with future interactions. **(C)** Harm sensitivity (*λ*) was higher for dictators with future interactions than those without future interactions. **(D)** For unfair splits, sensitivity to provocation completely mediated the relationship between interaction expectations and reactive punishment. **(E)** For fair splits, relative advantage completely mediated the relationship between interaction expectations and proactive punishment. *α*, sensitivity to provocation; *β*, relative advantage preference; *λ*, harm sensitivity; W, with future interactions; WO, without future interactions. \**p* < 0.05; \*\**p* < 0.01; \*\*\**p* < 0.001.

Moreover, the RSA analysis revealed a significant correlation between the self-reported closeness RDM and the punishment amounts RDM (*r* = 0.875, *p_permutation_* < 0.001, **Fig. 2E and Fig. S2**), further supporting the correspondence between subjective closeness and punishment behavior.

### Punishment of fair splits from dictators without future interactions depended on trial sequence

For the fair splits of dictators without future interactions, the ANOVA on the effects of trial sequence revealed a significant interaction between Preceding Split and Preceding Interaction expectation on punishment amounts (*F_1,29_* = 7.290, *p* = 1.144 × 10^−2^, *η^2^* = 0.201, **Fig. 2C**). Post-hoc comparisons showed that punishment of fair splits of dictators without future interactions was harsher when preceded by unfair than fair splits of dictators without future interactions (*p* = 7.640 × 10^−4^), but not influenced by fairness of splits from dictators with future interactions (*p* = 7.750 × 10^−1^). The effects of trial sequence were not significant in other conditions (all *p* > 0.05).

### The behavioral effects of interaction expectations corresponded to changes in sensitivity to provocation, relative advantage, and inflicted harm

The inequity aversion plus harm sensitivity model (M3, LOOIC = 5422.703, WAIC = 5350.903) had the lowest LOOIC and WAIC values compared to other models, including the standard inequity aversion model (M1, LOOIC = 6415.399, WAIC = 6341.068), the inequity aversion plus cost aversion model (M2, LOOIC = 5565.511, WAIC = 5501.916), and the inequity aversion plus retributive preference model (M4, LOOIC = 5631.247, WAIC = 5558.855). Thus, the inequity aversion plus harm sensitivity model was identified as the winning model. Moreover, both LOOIC and WAIC weights supported the superiority of M3 (both weights > 0.9999) over the competing models (both weights < 0.0001).

The winning model was further validated with two analyses. First, we observed a strong correlation between the model predictions and actual behavior across 30 participants (*r* = 0.884, *p =* 9.991×10^−11^, **Fig. S3A and 3B**) and across 160 trials (*r* = 0.941, *p =* 5.438×10^−76^, **Fig. S3C and 3D**). Moreover, the ANOVA on the simulated punishment amounts replicated the Split × Interaction expectation interaction observed in the actual data (*F_1,29_* = 28.352, *p* = 1.000×10^−5^, *η^2^* = 0.494, **Fig. S3E and 3F**). These results provided robust evidence that our model well predicted participants’ behaviors. Second, the parameter recovery analysis revealed a high correspondence between the true and recovered parameters (**Fig. S4**).

Analysis of the winning model’s estimated parameters revealed that interaction expectations influenced sensitivity to provocation (*α*), relative advantage (*β*), and inflicted harm (*λ*). In particular, participants exhibited higher sensitivity to provocation (95% HDI: [−0.573, −0.132], **Fig. 3A**), higher preferences for relative advantage (95% HDI: [0.057, 0.465], **Fig. 3B**), and lower sensitivity to inflicted harm (95% HDI: [0.095, 0.383], **Fig. 3C**) when interacting with dictators without future interactions than those with future interactions.

The validity of computational parameters was further examined with their associations with behavioral data. For unfair splits, sensitivity to provocation (*α*) acted as full mediators, completely mediating the effects of Interaction expectation on reactive punishment (indirect effect = 1.010, Bootstrap 95% CI [0.425, 1.969]; **Fig. 3D**), moreover, correlation analyses revealed no significant correlation between differences in inflicted harm (*λ*) and reactive punishment (*r* = −0.001, *p* > 0.05). For fair splits, sensitivity to relative advantage completely mediated the effects of Interaction expectation on proactive punishment (indirect effect = −0.557, Bootstrap 95% CI [−0.849, −0.283]; **Fig. 3E**). Moreover, correlation analyses indicated a significant correlation between differences in harm sensitivity (λ) and proactive punishment (*r* = −0.416, *p* = 2.236×10^−2^).

### The effects of interaction expectations on dmPFC and dlPFC responses were modulated by the fairness of splits

The interaction of Split × Interaction expectation revealed activation in the dmPFC and right dlPFC (**Table 1 and Fig. 4A**), with the latter identified using a more lenient cluster-defining threshold of *p* < 0.005. Post-hoc comparisons revealed that neural responses in these regions to unfair splits were stronger for dictators with future interactions than for those without. Conversely, their responses to fair splits were lower for dictators with future interactions compared to those without.

**Fig. 4.**
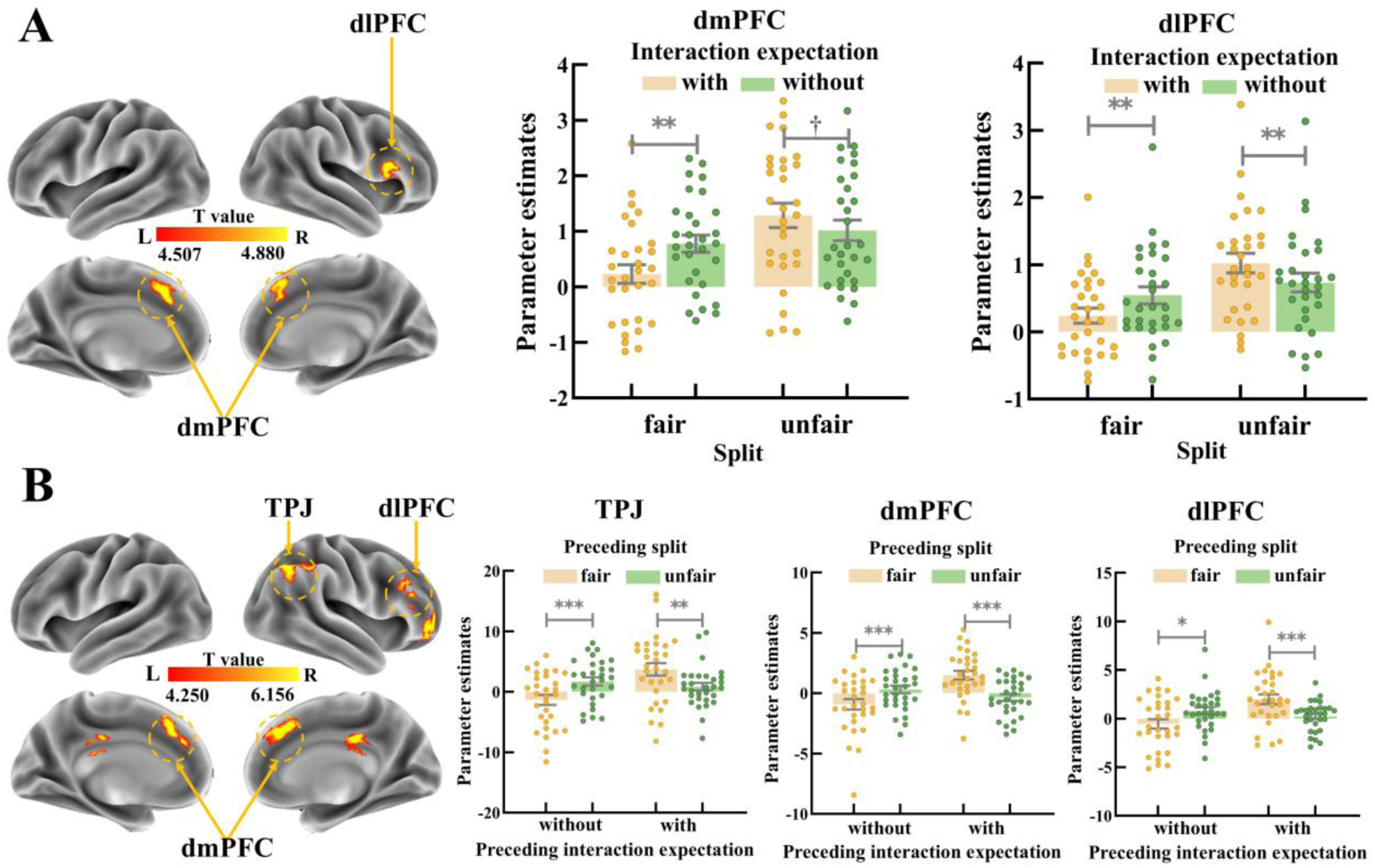
fMRI results. **(A)** Brain regions revealed by the interaction contrast between Interaction expectation and Split. The interaction revealed activation in the dmPFC and dlPFC. **(B)** Brain regions associated with the effects of trial sequence on punishment of fair splits from dictators without future interactions. The effect was associated activity in the TPJ, dmPFC, and dlPFC. dlPFC, dorsolateral prefrontal cortex; dmPFC, dorsomedial prefrontal cortex; TPJ, temporoparietal junction; L, left; R, right; with, with future interactions; without, without future interactions; ^†^*p* < 0.05, one-tailed; \**p* < 0.05; \*\**p* < 0.01; \*\*\**p* < 0.001.

**Table 1.** Brain regions identified in different analyses.

| Brain Regions | Cluster size (voxel) | L/R | MNI |  |  | Local Maximal T | Brodmann Area |
| --- | --- | --- | --- | --- | --- | --- | --- |
|  |  |  | Coordinates of Local Maxima (mm) |  |  |  |  |
|  |  |  | x | y | z |  |  |
|  |  |  | (with - without) unfair splits > (with - without) fair splits |  |  |  |  |
| Dorsomedial prefrontal cortex | 203 | L/R | -4 | 22 | 46 | 5.817 | BA 8/32 |
| Dorsolateral prefrontal cortex | 290 | R | 54 | 18 | 24 | 4.057 | BA 45 |
| **Preceding contexts: (unfair - fair)<sub>without</sub> > (unfair - fair)<sub>with</sub>** |  |  |  |  |  |  |  |
| Dorsomedial prefrontal cortex | 675 | L/R | 0 | 26 | 38 | 5.817 | BA 9/32 |
| Dorsolateral prefrontal cortex | 155 | R | 38 | 26 | 30 | 4.250 | BA 46 |
| Temporoparietal junction | 542 | R | 48 | -58 | 42 | 5.122 | BA 40 |
| Middle cingulate cortex | 149 | L/R | 6 | -28 | 32 | 6.156 | BA 23 |
| Ventrolateral prefrontal cortex | 247 | R | 36 | 60 | 2 | 4.629 | BA 10 |
| **Subjective utility** |  |  |  |  |  |  |  |
| Ventromedial prefrontal cortex | 215 | L/R | -8 | 44 | -12 | 4.426 | BA 11 |
| Precuneus | 169 | L | -6 | -50 | 62 | 5.439 | BA 5 |
| Superior occipital gyrus | 381 | L | -16 | -98 | 20 | 5.391 | BA 19 |
| **Sensitivity to provocation ($\alpha$)** |  |  |  |  |  |  |  |
| Dorsal anterior cingulate cortex | 252 | L/R | -4 | 26 | 18 | 7.006 | BA 24 |
| **Relative advantage ($\beta$)** |  |  |  |  |  |  |  |
| Dorsolateral prefrontal cortex | 149 | R | 24 | 22 | 54 | -5.118 | BA 8 |
| **Harm sensitivity ($\lambda$)** |  |  |  |  |  |  |  |
| Temporoparietal junction | 289 | R | 38 | -46 | 30 | -4.077 | BA 40 |

L, left; R, right.

### Neural signatures underlying trial sequence effects on punishment of fair splits from dictators without future interactions

For the fair splits from dictators without future interactions, the interaction between Preceding Interaction expectation (with/without) and Preceding Split (fair/unfair) identified the right TPJ, dlPFC, ventrolateral prefrontal cortex, bilateral dmPFC, and middle cingulate cortex (**Table 1 and Fig. 4B**). These regions showed greater activity for trials following unfair than fair splits by dictators without future interactions, whereas they show greater activity for trials following fair than unfair splits by dictators with future interactions.

### Neural responses tracking subjective utility of punishment decision-making

The parametric modulations of subjective utility revealed the bilateral ventromedial prefrontal cortex, left precuneus, and superior occipital gyrus, indicating their role in encoding the subjective utility of punishment decisions (**Table 1 and Fig. 5A**).

**Fig. 5.**
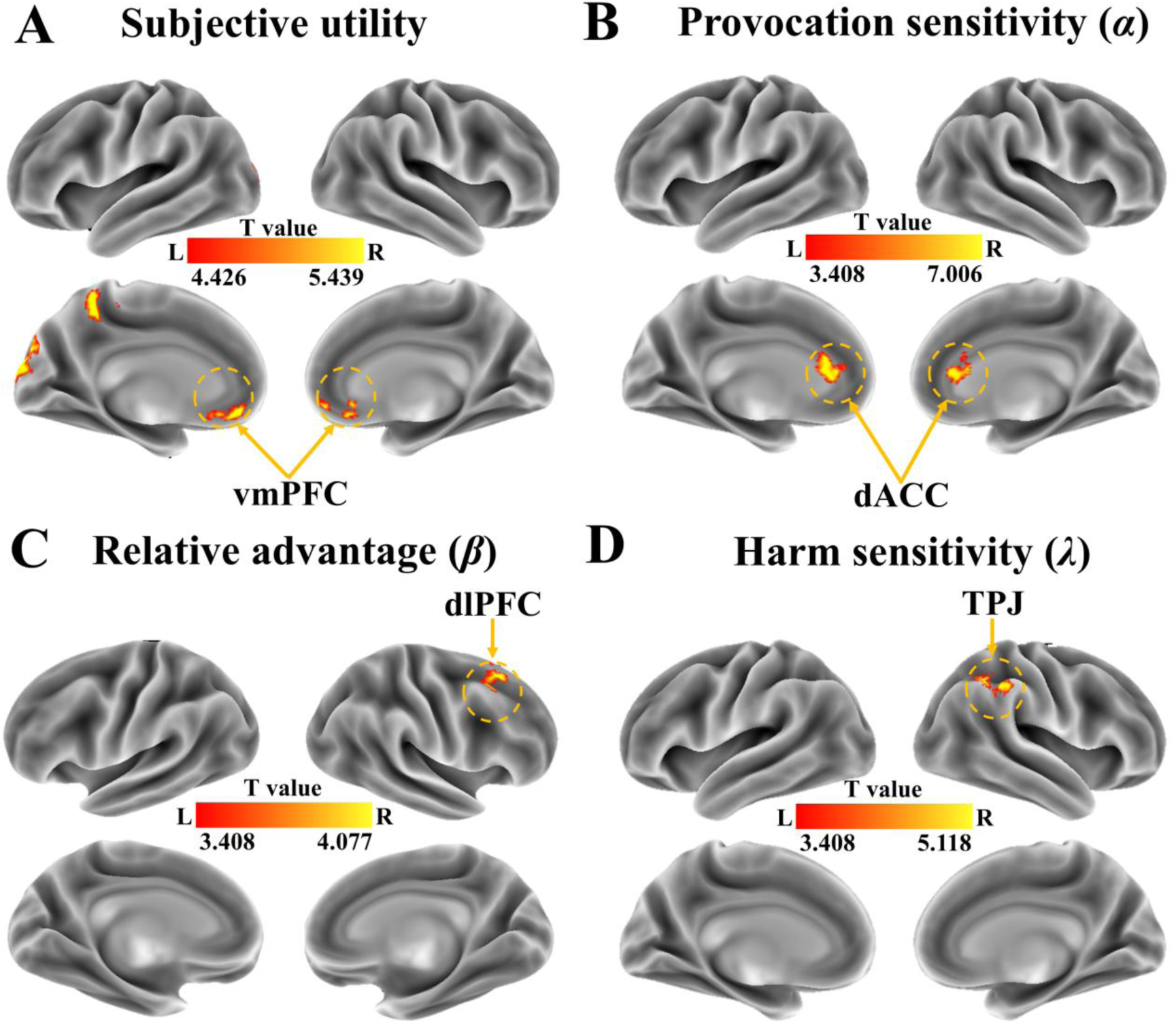
Brain responses modulated by subjective utility and parameter values. **(A)** Brain regions exhibiting associations with subjective utility. **(B)** Brain regions associated with changes in sensitivity to provocation (*α*) under unfair splits (without vs. with). **(C)** Brain regions associated with the changes in the relative advantage preferences (*β*) under fair splits (with vs.without) **(D)** Brain region associated with changes in harm sensitivity (*λ*) under fair splits (with vs.without). vmPFC, ventromedial prefrontal cortex; dACC, dorsal anterior cingulate cortex; dlPFC, dorsolateral prefrontal cortex; TPJ, temporoparietal junction; L, left; R, right.

### Brain responses modulated by sensitivity to norm violations, relative advantage, and inflicted harm

To further investigate the neural signatures associated with the impact of interaction expectations on different components of subjective utility, we examined brain responses modulated by individual differences in the effects of interaction expectations on sensitivity to provocation (*α*), relative advantage (*β*), and inflicted harm (*λ*). The results revealed a positive association between changes in values of parameter *α* and dACC responses to unfair splits (**Table 1 and Fig. 5B**). Specifically, larger dACC responses to unfair splits from dictators without (vs. with) future interactions were associated with a greater sensitivity to provocation committed by dictators without future interactions. Moreover, a negative association was found between changes in values of parameter *β* and right dlPFC responses to fair splits (**Table 1 and Fig. 5C**), indicating that larger dlPFC responses to fair splits from dictators without (vs. with) future interactions were linked to stronger preferences for relative advantage over dictators without future interactions. Lastly, we observed a negative association between changes in values of parameter *λ* and right TPJ responses to fair splits (with cluster-defining threshold of *p* < 0.005, **Table 1 and Fig. 5D**), such that larger TPJ responses to fair splits from dictators without (vs. with) future interactions were associated with lower harm sensitivity for dictators without future interactions.

## Discussion

Our findings demonstrate that future interaction prospects modulate both reactive and proactive costly punishment. Participants imposed harsher punishment on both fair and unfair behaviors committed by individuals they did not expect to interact with, a pattern that corresponded with lower self-reported social closeness. Heightened sensitivity to norm violations, as reflected in dACC activity, drove increased reactive punishment toward wrongdoers without future interactions. In contrast, punishment of future-interacting transgressors was associated with enhanced dlPFC and dmPFC activity. A stronger preference for relative advantage and diminished harm sensitivity, respectively linked to dlPFC and TPJ activity, mediated higher proactive punishment toward non-future-interacting individuals. Notably, proactive punishment and its underlying neurocognitive processes followed a collective retaliation pattern, increasing after unfair (vs. fair) behavior from another non-future-interacting individual. Collectively, these findings provide a mechanistic account of how future interaction prospects shape reactive and proactive costly punishment.

Despite extensive research, findings on the relationship between social bonds and reactive punishment remain mixed, likely due to the multidimensional nature of social bonds [12, 21]. Our study clarifies this by showing that future interaction prospects—a fundamental dimension of social bonds—modulate reactive punishment. Consistent with the mere preference hypothesis [12], participants imposed harsher punishment on norm violators without future interactions, whom they perceived as less close. This effect stemmed from heightened sensitivity to provocation, suggesting a mediating role of moral outrage. Supporting this, neuroimaging results revealed that greater provocation sensitivity correlated with increased dACC activity toward transgressors without future interactions. The dACC is consistently implicated in encoding salience and aversive reactions to norm violations [41, 60], aligning with its broader role in processing negative emotions such as pain and disgust [61]. For instance, dACC responses to provocation amplify under aversive emotional states [62] and diminish with cognitive reappraisal [35]. Thus, our computational and neuroimaging findings suggest that harsher reactive punishment for transgressors without future interactions reflects intuitive, emotionally driven responses shaped by affective preferences.

In contrast, participants were more lenient toward norm violators with future interaction prospects, corresponding to increased dmPFC and dlPFC activity. These regions integrate diverse sources of information (e.g., intentions and contextual cues) to guide flexible, context-dependent punishment [3, 35, 39, 63]. Thus, reactive punishment of future-interacting transgressors may reflect a deliberate strategy for norm enforcement rather than a reflexive expression of outrage. In accordance, participants punished future-interacting transgressors as often as those without future interactions (see Supplementary Results) but imposed milder sanctions, likely serving as warnings to promote behavioral correction [64]. Similarly, reactive punishment of in-group perpetrators has been linked to self-reported motives to encourage normative adherence [65]. These findings delineate distinct motives underlying reactive punishment across social contexts, reconciling the tension between norm enforcement and affective preference accounts.

Beyond reactive punishment, participants also exhibited stronger proactive punishment toward fair offers from individuals without future interactions. Computational modeling indicated that this behavior was driven by changes in both relative advantage concerns and harm sensitivity. On the one hand, the motive to gain advantage through proactive punishment aligns with behavioral economic findings showing that proactive punishment is rare when it does not confer relative advantage—for example, when cost-to-impact ratios are low or when only nonmonetary sanctions are available [33, 66]. Furthermore, social psychology research on self-enhancement consistently demonstrates that individuals employ behavioral strategies and cognitive distortions to maintain a sense of superiority for themselves and close others over distant others [32]. Our findings integrate these perspectives by providing novel evidence that the pursuit of relative advantage is amplified in interactions with distant others, even at economic and moral costs. The status-seeking motive was reflected in dlPFC activity, consistent with its established role in status-related decision-making and learning [67]. These results suggest that proactive punishment as a means of status-seeking involves strategic considerations, requiring greater attentional resources and extensive integration of information, processes supported by the dlPFC.

On the other hand, the modulation of proactive punishment was linked to lower harm sensitivity, which correlated with TPJ activity. Consistent with our findings, the TPJ has been causally implicated in proactive punishment [68]. However, our results provide novel evidence specifying the computations implemented in this region—namely, harm sensitivity [47, 69]. The observed changes in harm sensitivity and associated TPJ activity likely reflect a collective retaliation motive. Specifically, individuals tend to depersonalize distant others, perceiving them as interchangeable [29], thereby reducing sensitivity to harming an innocent individual for the wrongdoing of another within the same category [70]. Thus, the TPJ may facilitate moral justification processes, enabling individuals to rationalize collective punishment by constructing narratives that attribute collective responsibility to an uninvolved individual based on his social association with the actual offender [31]. Supporting this interpretation, proactive punishment and TPJ activity exhibited a collective retaliation pattern, enhancing after unfair (vs. fair) offers from another non-future-interacting individual. In short, our findings reveal that proactive punishment toward distant others is driven by both status-seeking and collective retaliation motives.

Several limitations merit consideration. First, although the current sample size aligns with prior neuroimaging studies [41], replication in larger, more diverse cohorts is needed to validate our findings. Second, while debates continue on whether second- and third-party punishment rely on distinct mechanisms [41, 42], our study revealed converging results across behavioral, computational, and neuroimaging levels for both types of punishment. Determining the boundary conditions that differentiate these forms of punishment lies beyond the scope of this study and warrants future investigation. Third, although brain–behavior correlations and previous literature support our interpretations of the implicated neural regions, the fMRI findings indicate regional involvement rather than causality. Elucidating the precise functions of these regions will require more targeted methodologies. Fourth, future research should examine additional dimensions of social bonds, such as similarity and interdependence, to provide a more comprehensive understanding of how social bonds modulate punitive decision-making. Finally, although proactive punishment is prevalent in both experimental contexts and real-world interactions [9, 71], socially desirable concerns may suppress such behaviors. Future investigations employing more sensitive and ecologically valid designs are needed to better capture this fundamental aspect of human social behavior.

Together, our study provides a unified framework for the neurocomputational mechanisms linking future interaction prospects to both reactive and proactive punishment. These findings advance our understanding of the motivational, cognitive, and emotional processes underlying costly punishment within social bonds, highlighting the interplay of multiple neuropsychological subcomponents, including provocation sensitivity, concerns for relative advantage, and harm sensitivity.

## Supporting information

Supplemental results and figures

## Acknowledgements

This study was supported by the National Natural Science Foundation of China (32271126, 31920103009), grant from Research Center for Brain Cognition and Human Development, Guangdong, China (No. 2024B0303390003), and grant from Striving for the First-Class, Improving Weak Links and Highlighting Features (SIH) Key Discipline for Psychology in South China Normal University, the Major Project of National Social Science Foundation (20&ZD153), and Shenzhen-Hong Kong Institute of Brain Science – Shenzhen Fundamental Research Institutions (2019SHIBS0003).

