## Supplemental results and figures for "Neurocomputational Mechanisms Linking Future Interaction Prospects to Reactive and Proactive Costly Punishment"

**Supplementary Materials**

**Supplementary Results**

**Rates of punishment.** The ANOVA on rates of punishment revealed significant main effects of Split (*F_1,29_* = 288.897, *p* = 1.282 × 10^-16^, *η^2^_partial_* = 0.909) and Interaction expectation (*F_1,29_* = 9.023, *p* = 5.447 × 10^−3^, *η^2^_partial_* = 0.237), indicating that participants punished more frequently to unfair splits than fair splits and dictators without future interactions than those with future interactions. In addition, the interaction of Split × Interaction expectation was significant (*F_1,29_* = 4.924, *p* = 3.446 × 10^−2^, *η^2^_partial_* = 0.145, **Fig. S1**). Post-hoc comparisons indicated that participants exhibited more frequent punishment for fair splits from dictators without future interactions (*p* = 8.2 × 10^−5^). For unfair splits, the punishment rates were not significantly different between dictators with and without future interactions (*p* = 4.354 × 10^−1^).

**Supplementary Figures**

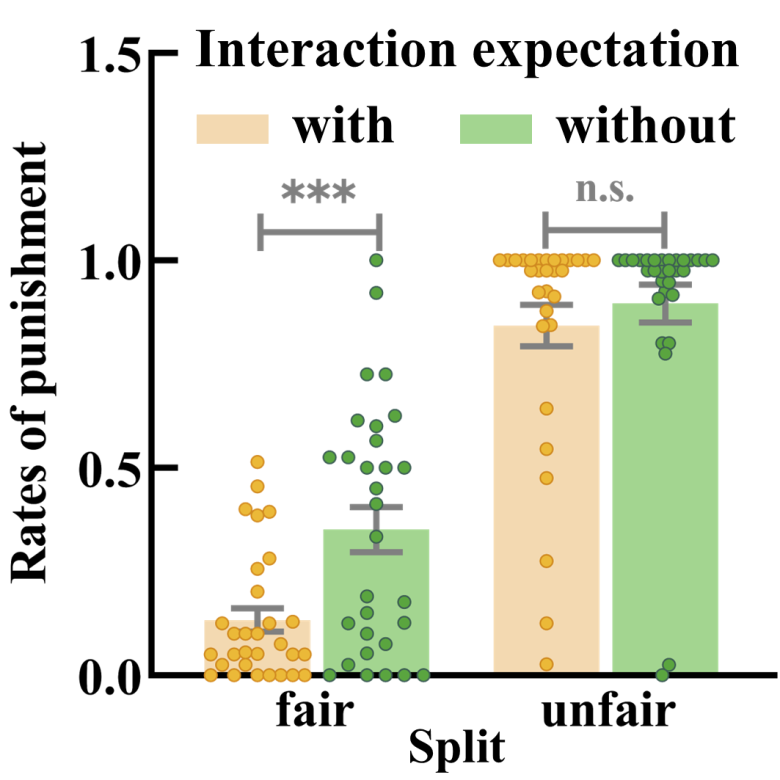

**Fig. S1 | Rates of punishment.** Participants punished fair splits of dictators without future interactions more frequently than those from dictators with future interactions, while punishment rates were comparable between unfair splits of dictators with and without future interactions. With, with future interactions; Without, without future interactions; ****p* < 0.001; n.s., not significant.

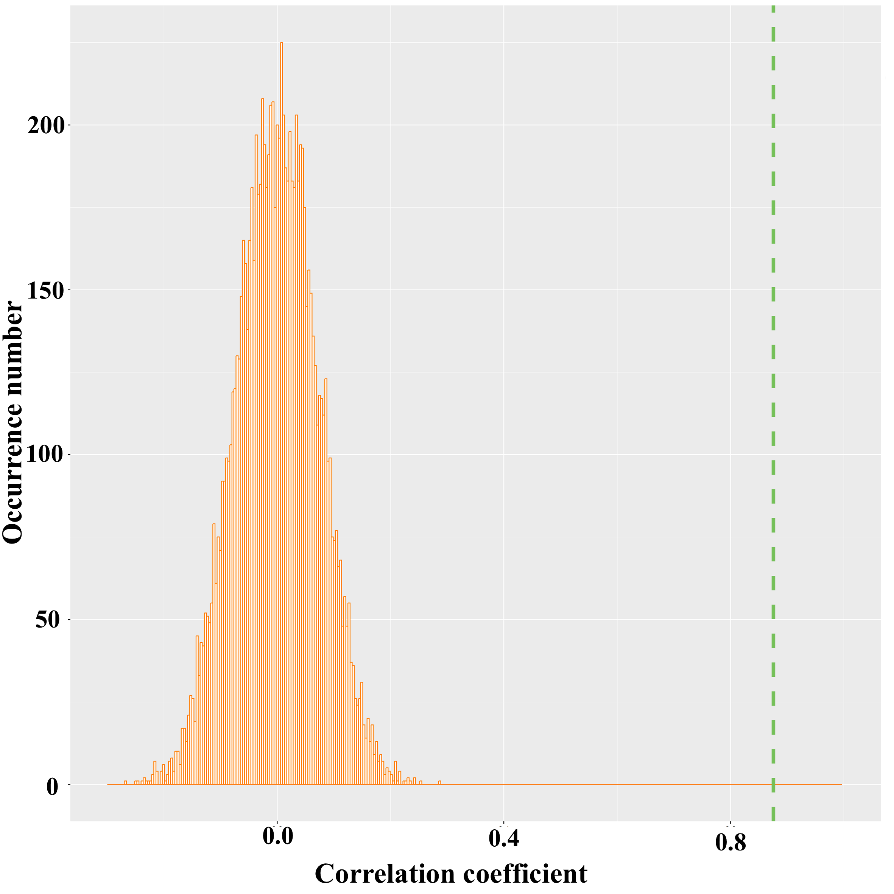

**Fig. S2 |** **Permutation test of RSA analysis on self-reported closeness and punishment amounts.** Permutation distribution of the correlation coefficient. The value obtained using real scores is indicated by the green dashed line.

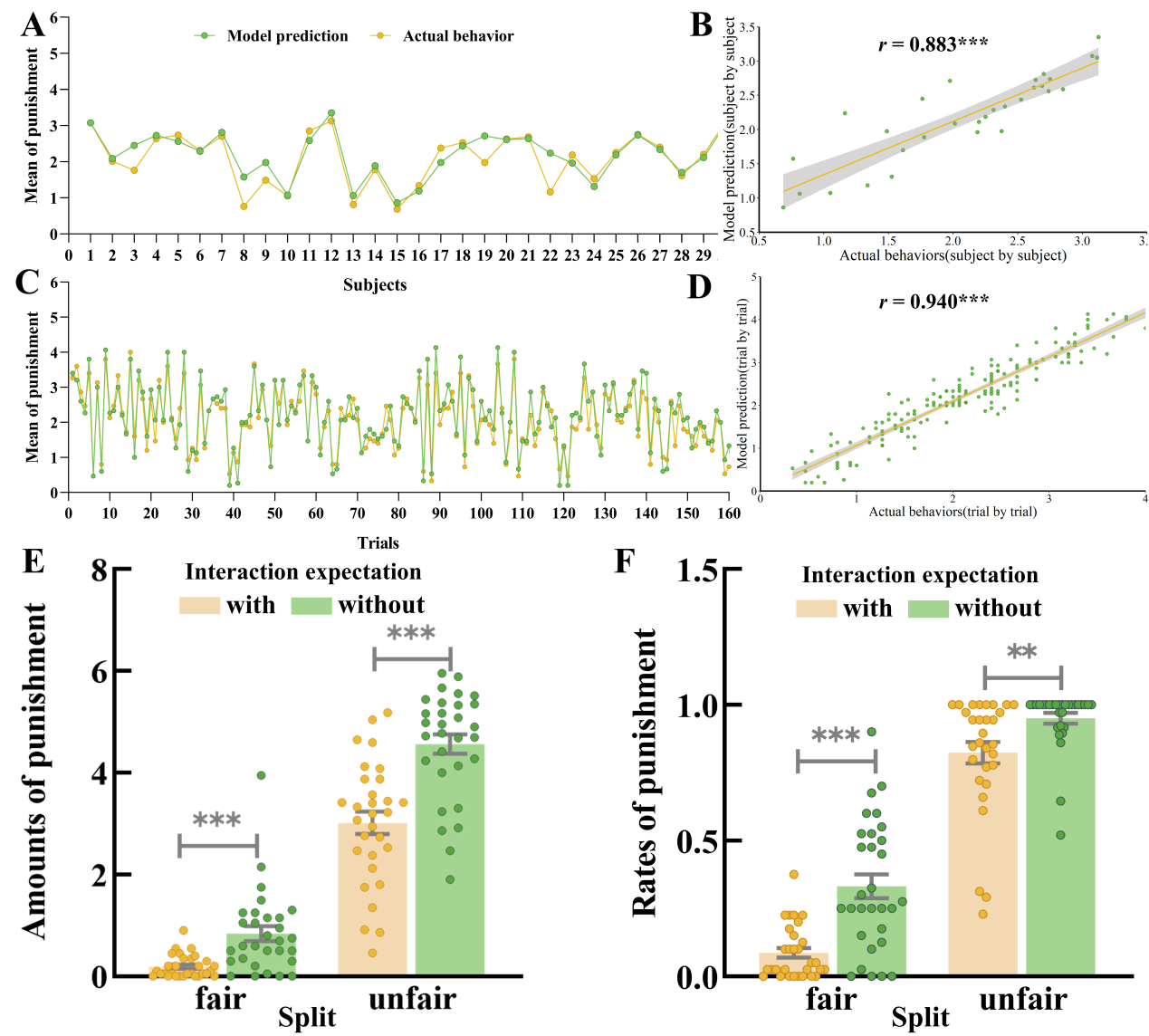

**Fig. S3 | Model validation. (A)** Consistency between actual behaviors and model predictions in the average level of costly punishment across 30 participants. **(B)** Correlation between actual behaviors and model predictions in the mean level of costly punishment across participants. **(C)** Consistency between actual behaviors and model predictions in the average level of costly punishment across 160 trials. **(D)** Correlation between actual behaviors and model predictions in the mean level of costly punishment across trials. **(E)** Average amounts of punishment predicted by the winning model. **(F)** Average rates of punishment predicted by the winning model. With, with future interactions; Without, without future interactions; ***p* < 0.01; ****p* < 0.001.

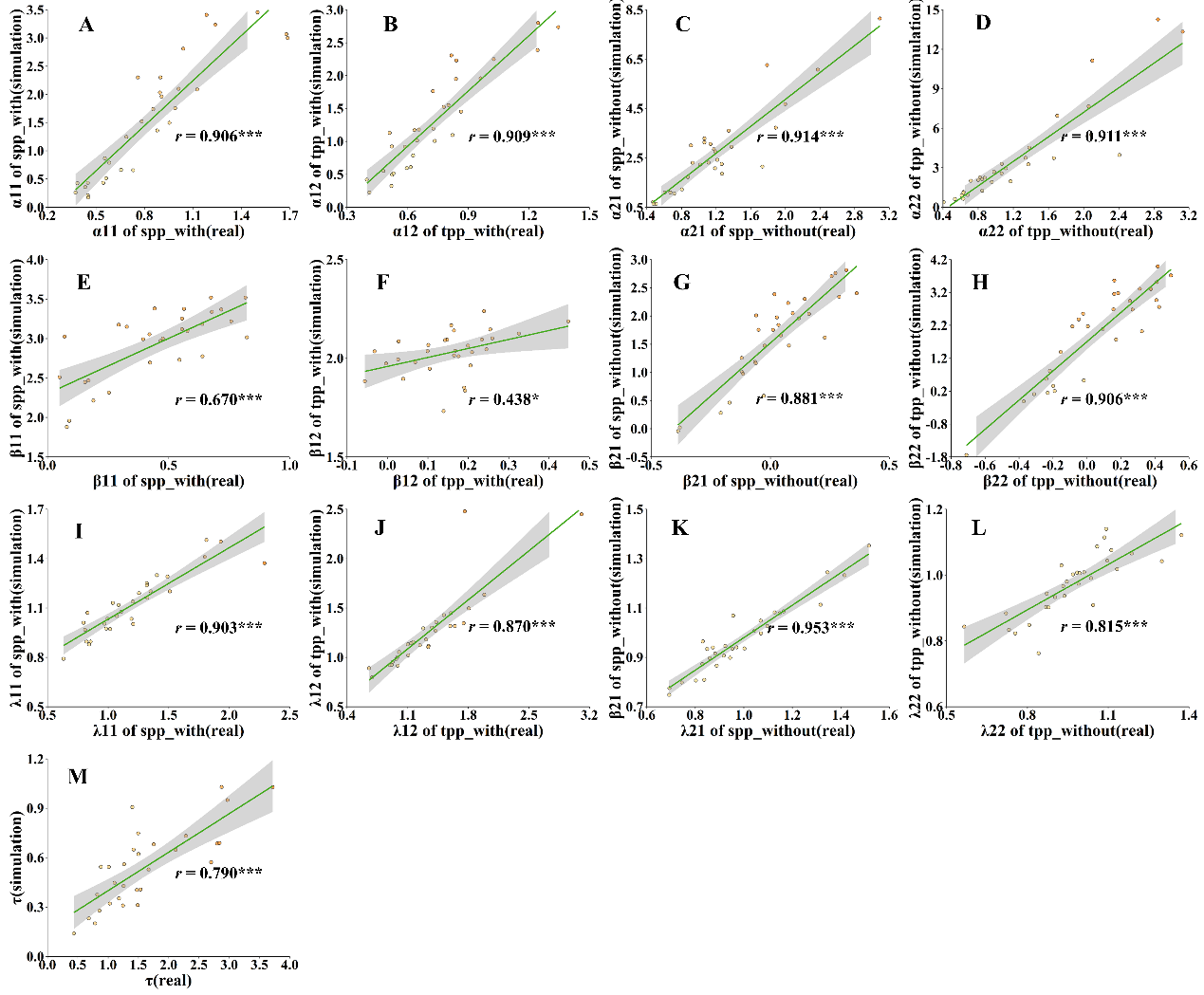

**Fig. S4 | Parameter recovery. (A)** Correlation between actual and simulated parameter values of sensitivity to provocation (α11) for dictators with future interactions in the second-party role. **(B)** Correlation between actual and simulated parameter values of sensitivity to provocation (α12) for dictators with future interactions (α12) in the third-party role. **(C)** Correlation between actual and simulated parameter values of sensitivity to provocation (α21) for dictators without future interactions in the second-party role. **(D)** Correlation between actual and simulated parameter values of sensitivity to provocation (α22) for dictators without future interactions in the third-party role. **(E)** Correlation between actual and simulated parameter values of relative advantage (β11) for dictators with future interactions in the second-party role**. (F)** Correlation between actual and simulated parameter values of relative advantage (β12) for dictators with future interactions in the third-party role. **(G)** Correlation between actual and simulated parameter values of relative advantage (β21) for dictators without future interactions in the second-party role. **(H)** Correlation between actual and simulated parameter values of relative advantage (β22) for dictators without future interactions in the third-party role. **(I)** Correlation between actual and simulated parameter values of inflicted harm (λ11) for dictators with future interactions in the second-party role. **(G)** Correlation between actual and simulated parameter values of inflicted harm (λ12) for dictators with future interactions in the third-party role. **(K)** Correlation between actual and simulated parameter values of inflicted harm (λ21) for dictators without future interactions in the second-party role. **(L)** Correlation between actual and simulated parameter values of inflicted harm (λ22) for dictators without future interactions in the third-party role. **(M)** Correlation between actual and simulated parameter values of τ. The colors of dots indicated the differences between actual behaviors (x-axis) and model predictions (y-axis) from the largest (brown) to the smallest (yellow). With, with future interactions; Without, without future interactions; SPP, second-party punishment; TPP, third-party punishment; **p* < 0.05; ****P* < 0.0005.
